# The mutational landscape of a prion-like domain

**DOI:** 10.1101/592121

**Authors:** Benedetta Bolognesi, Andre J. Faure, Mireia Seuma, Jörn M. Schmiedel, Gian Gaetano Tartaglia, Ben Lehner

**Author notes:** These authors contributed equally to this work (Benedetta Bolognesi, Andre J. Faure).

## Abstract

Specific insoluble protein aggregates are the hallmarks of many neurodegenerative diseases^1–5^. For example, cytoplasmic aggregates of the RNA-binding protein TDP-43 are observed in 97% of cases of Amyotrophic Lateral Sclerosis (ALS)^6,7^. However, it is still unclear for ALS and other diseases whether it is the insoluble aggregates or other forms of the mutated proteins that cause these diseases that are actually toxic to cells^8–13^. Here we address this question for TDP-43 by systematically mutating^14^ the protein and quantifying the effects on cellular toxicity. We generated >50,000 mutations in the intrinsically disordered prion-like domain (PRD) and observed that changes in hydrophobicity and aggregation potential are highly predictive of changes in toxicity. Surprisingly, however, increased hydrophobicity and cytoplasmic aggregation actually reduce cellular toxicity. Mutations have their strongest effects in a central region of the PRD, with variants that increase toxicity promoting the formation of more dynamic liquid-like condensates. The genetic interactions in double mutants reveal that specific structures exist in this ‘unstructured’ region *in vivo*. Our results demonstrate that deep mutagenesis is a powerful approach for probing the sequence-function relationships of intrinsically disordered proteins as well as their *in vivo* structural conformations. Moreover, we show that aggregation of TDP-43 is not harmful but actually protects cells, most likely by titrating the protein away from a toxic liquid-like phase.

## Main text

The conversion of specific proteins into insoluble aggregates is a hallmark of many neurodegenerative disorders, including Alzheimer’s, Parkinson’s, Huntington’s, and Amyotrophic Lateral Sclerosis (ALS) with dominantly inherited mutations in aggregate-forming proteins causing rare familial forms of these diseases^3,6,15^. However, both in humans and in animal models, there is often only a weak association between the presence of aggregates and disease progression^16,17^. Indeed, many therapeutic approaches that reduce the formation of aggregates have failed at different stages of development^12,18,19^. On the other hand, there is increasing evidence that alternative protein assemblies generated during or in parallel to the aggregation process, may be toxic^8–11,20^. Despite evidence that cellular damage may be induced either before, after or independent of the formation of insoluble aggregates, the latter are still widely assumed to be pathogenic in many neurodegenerative diseases^21,22^.

For many proteins, aggregation depends critically on intrinsically disordered regions with a low sequence complexity resembling that of infectious yeast prions. These prion-like domains (PRDs) are also enriched in proteins that can form liquid-like cellular condensates^23–25^ with the PRDs necessary and sufficient for liquid-demixing^26,27^. At least *in vitro*, insoluble aggregates can nucleate from more liquid phases^27–29^, leading to the suggestion that liquid de-mixed states can mature into pathological aggregates^22^. Disordered regions^30^ and low complexity sequences^31^ are also enriched in dosage-sensitive proteins – those that are toxic when their concentration is increased. At least for one model protein that has been tested, however, it is the formation of a concentration-dependent liquid-like phase – not aggregation – that causes cellular toxicity^31^. Similarly, the toxicity of two mutant forms of the prion Sup35 could be explained only on the basis of their ability to populate a non-aggregate, liquid-like state^23,32^.

Cytoplasmic aggregates of the RNA-binding protein TDP-43 are a hallmark of ALS, present in 97% of post-mortem samples^6,7^. TDP-43 aggregates are also present at autopsy in nearly all cases of frontotemporal dementia (FTD) that lack tau-containing inclusions (about half of all cases of FTD which is the second most common dementia)^33^. TDP-43 aggregates are also a hallmark of inclusion body myopathy and a secondary pathology in Alzheimer’s, Parkinson’s, and Huntington’s disease^33–35^. However, TDP-43 aggregates are also observed – albeit at low frequency – in control samples^36^ and, *in vitro*, TDP-43 can form both amyloid aggregates and liquid condensates^37–41^. Mutations in TDP-43 cause ∼5% of familial ALS cases^17,42^, with these mutations clustered in the PRD and reported to interfere with nuclear-cytoplasmic transport, RNA processing, splicing, and protein translation^16,43–48^. However, despite extensive investigation, the molecular form of the protein that causes cellular toxicity is still unknown^16,49^.

We reasoned that systematic (‘deep’) mutagenesis could be an unbiased approach to identify and investigate the toxic species of proteins^14,50,51^. A map of which amino acid changes increase or decrease the toxicity of a protein to a cell should, if sufficiently comprehensive, clarify both the properties of the protein and its *in vivo* structural conformations associated with toxicity^52^. The effects of a small number of mutations on TDP-43 toxicity or aggregation have been previously reported^10,37,53–56^. However, on the basis of a handful of mutations, the relationship between aggregation and toxicity is far from clear.

We used error-prone oligonucleotide synthesis to comprehensively mutate the PRD of TDP-43. We introduced the library into yeast cells, induced expression and used deep sequencing before and after induction to quantify the relative effects of each variant on growth in three biological replicates (Fig. 1a). After quality control and filtering (ED Fig. 1a-c), the dataset quantifies the relative toxicity of 1,266 single and 56,730 double amino acid (AA) changes in the PRD with high reproducibility (Fig. 1b, ED Fig. 1d,e). The toxicity scores also correlate very well with the toxicity of the same variants re-tested in the absence of competition (Fig. 1c).

**Figure 1.**
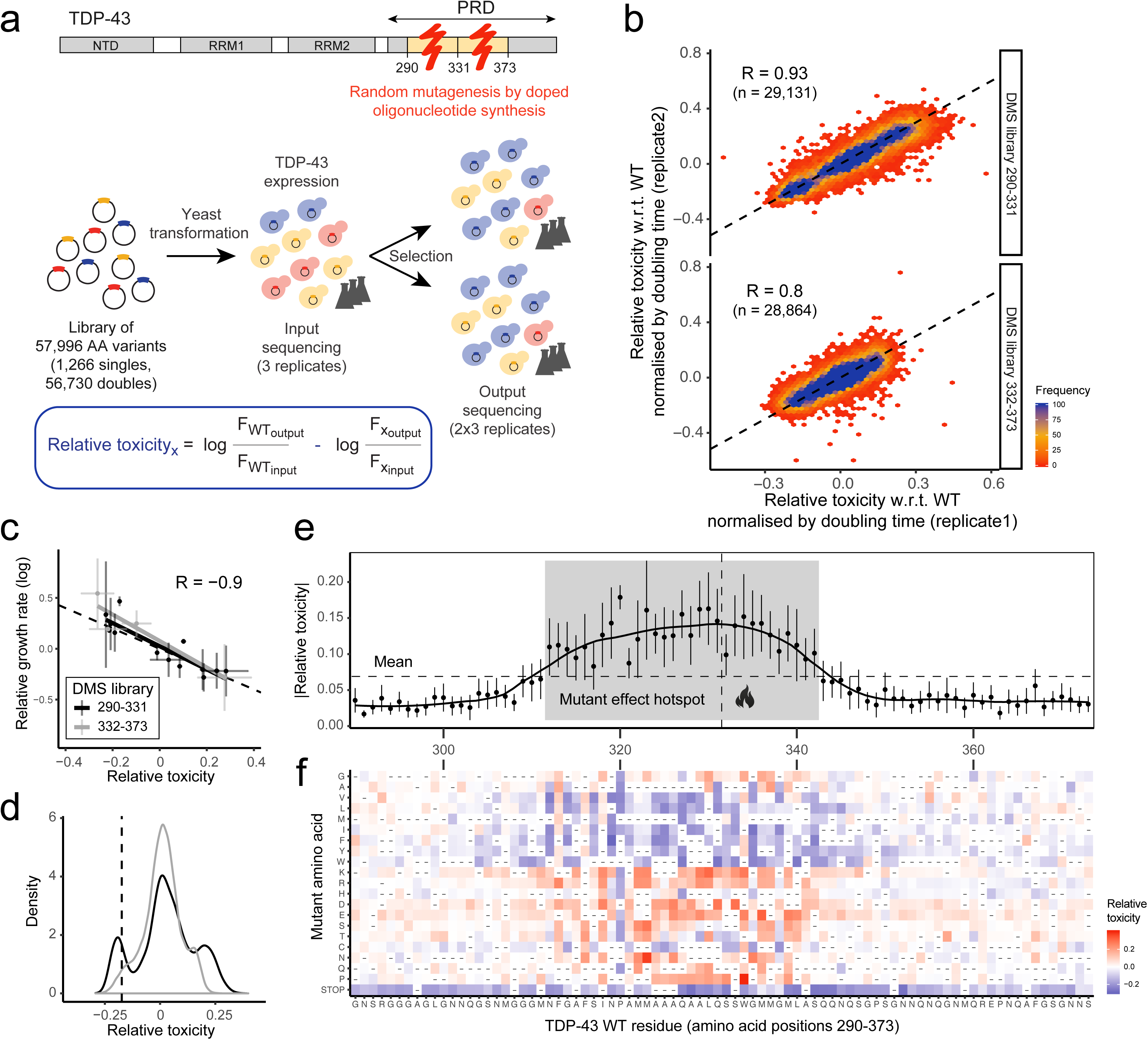
Deep mutational scanning (DMS) of the prion-like domain (PRD) of TDP-43. **a** Domain structure of TDP-43 and DMS experimental protocol: For each library, 3 independent selection experiments were performed. In each experiment one input culture was split into two cultures for selection upon induction of TDP-43 expression (6 outputs total). Relative toxicity of variants was calculated from changes of output to input frequencies relative to WT. **b** Correlation of toxicity estimates between replicates 1 and 2 for single and double amino acid (AA) mutants shown separately for each library (290-332; 332-373). The Pearson correlation coefficients (R) are indicated. Toxicity correlations between all replicates are shown in ED Fig. 1d, e. **c** Comparison of toxicity from pooled selections and individually measured growth rates for selected variants. Vertical and horizontal error bars indicate 95% confidence intervals of mean growth rates and toxicity estimates respectively. Linear fits of the data are shown separately for each library and Pearson correlation (R) after pooling data from both libraries is indicated. **d** Toxicity distribution of single and double mutants, shown separately for each library (colour key as in panel c). WT variant has toxicity of zero, mean toxicity of variants with single STOP codon mutation is indicated by dashed vertical line. **e** Absolute toxicity of single mutants stratified by position. Error bars indicate 95% confidence intervals of mean (per-position) toxicity estimates. A local polynomial regression (loess) over toxicity estimates of all single mutants is shown. The vertical dashed line indicates the boundary between the two DMS libraries. The horizontal dashed line indicates the mean absolute toxicity of all single mutants. The mutant effect “hotspot” (mean per-position |toxicity| > mean |toxicity|) is highlighted in grey. **f** Heatmap showing single mutant toxicity estimates. The vertical axis indicates the identity of the substituted (mutant) AA. Heatmap cells of variants not present in the library are denoted by “-“.

The toxicity of both single and double mutants had a tri-modal distribution (Fig. 1d, ED Fig. 2a-c), with 18,023 variants more toxic and 16,152 variants less toxic than wild-type (WT) TDP-43 (t-test false discovery rate, FDR=0.05). The dataset therefore allows us to investigate how mutations both increase and decrease toxicity. Very interestingly, all recurrent familial ALS (fALS) TDP-43 mutations increase toxicity, with a strong bias towards moderate effects (t-test p-value = 0.016) (ED Fig. 2d).

**Figure 2.**
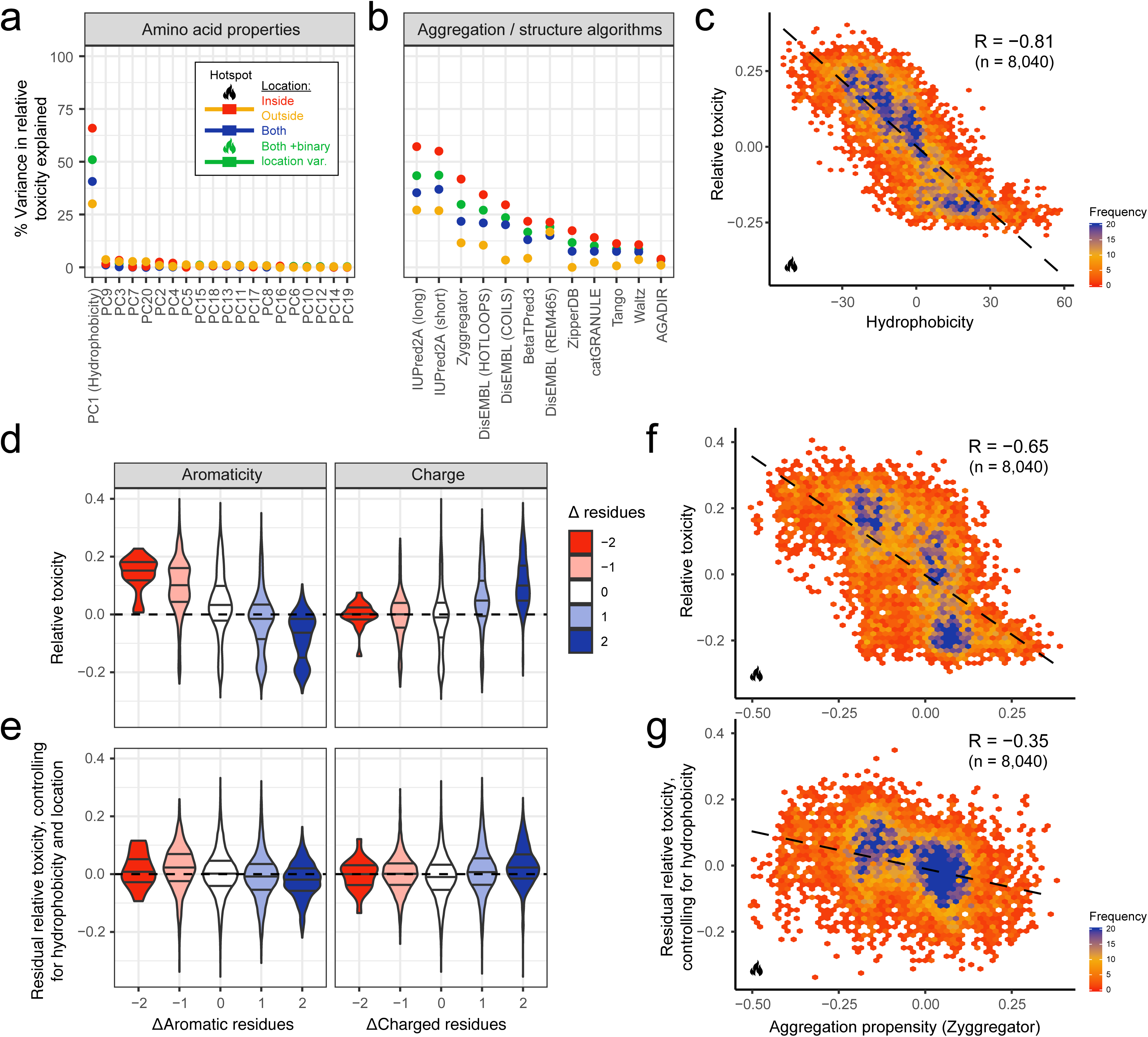
Changes in hydrophobicity are highly predictive of TDP-43 cellular toxicity. **a** Percentage variance of toxicity explained by linear regression models predicting single and double mutant variant toxicity from changes in AA properties upon mutation (PCs, principal components of a collection of AA physico-chemical properties). Different regression models were built for different subsets of the data. Simple linear regression models for all variants (blue) or only variants inside (red) or outside (yellow) the hotspot region. And a regression model using all variants and including a binary location variable (inside/outside hotspot) as well as an interaction term between binary location variable and the indicated AA property feature (green). **b** Percentage variance of toxicity explained by linear regression models predicting variant toxicity using scores from aggregation/structure algorithms (see Methods). Colour key shown in panel a. See also ED Fig. 4. **c** Toxicity of variants with single or double mutations within the hotspot region as a function of hydrophobicity changes (PC1) induced by mutation. The Pearson correlation (R) before binning is indicated. See also ED Fig. 7a. **d** Toxicity distributions of single and double mutants stratified by the change in the number of aromatic (H,F,W,Y,V) or charged residues (R,D,E,K) relative to the WT sequence. Horizontal axis as in panel e. **e** Distribution of residual toxicity after controlling for the effect of hydrophobicity and location on toxicity (green regression model in panel a) stratified by the number of aromatic (H,F,W,Y,V) or charged (R,D,E,K) AAs. **f** Single and double mutant variant toxicity as a function of changes in aggregation propensity (Zyggregator). Only variants occurring within the toxicity hotspot are depicted. The Pearson correlation (R) before binning is indicated. **g** Toxicity as a function of aggregation propensity after controlling for hydrophobicity (red regression model in panel a). Only variants occurring within the toxicity hotspot are depicted. The Pearson correlation (R) before binning is indicated. See also ED Fig. 7b.

Plotting the mean toxicity of all mutations at each position in the sequence reveals a 31 AA hotspot (312-342) where the effects of mutations are strongest (Fig. 1e). The variance in toxicity per position is also the highest within this hotspot, with mutations both strongly increasing and decreasing toxicity (Fig. 1e). A heatmap of the toxicity of all of the single mutations also clearly reveals this hotspot, with most mutations of strong positive or negative effect falling within this 31 AA window (Fig. 1f). Equally strikingly, mutations to the same AA but in different positions within the hotspot often have very similar effects (Fig. 1f). In particular, mutations to charged and polar residues increase toxicity throughout the hotspot and mutations to hydrophobic amino acids decrease toxicity (Fig. 1f).

To more systematically identify features associated with changes in toxicity we made use of all 53,468 variants carrying one or two AA substitutions. We used principal components analysis (PCA) to reduce the redundancy in a list of over 350 AA physicochemical properties (ED Fig. 3) and linear regression to quantify how well changes in these physicochemical properties predict changes in the toxicity of TDP-43. A principal component very strongly related to hydrophobicity is the most predictive feature of toxicity, explaining 66% of the variance in toxicity of all 8,040 mutants within the 312-342 hotspot and 51% of the variance in toxicity of all genotypes (Fig. 2a). With the same approach we tested the performance of established predictors of protein aggregation, intrinsic disorder and other properties. None of them are as predictive as hydrophobicity (Fig. 2b, c). Importantly, after controlling for hydrophobicity, additional features such as charge and aromaticity do not predict toxicity (Fig. 2d, e, ED Fig. 4a) with aggregation potential accounting for an additional 4% of variance in the hotspot (Fig. 2f, g).

**Figure 3.**
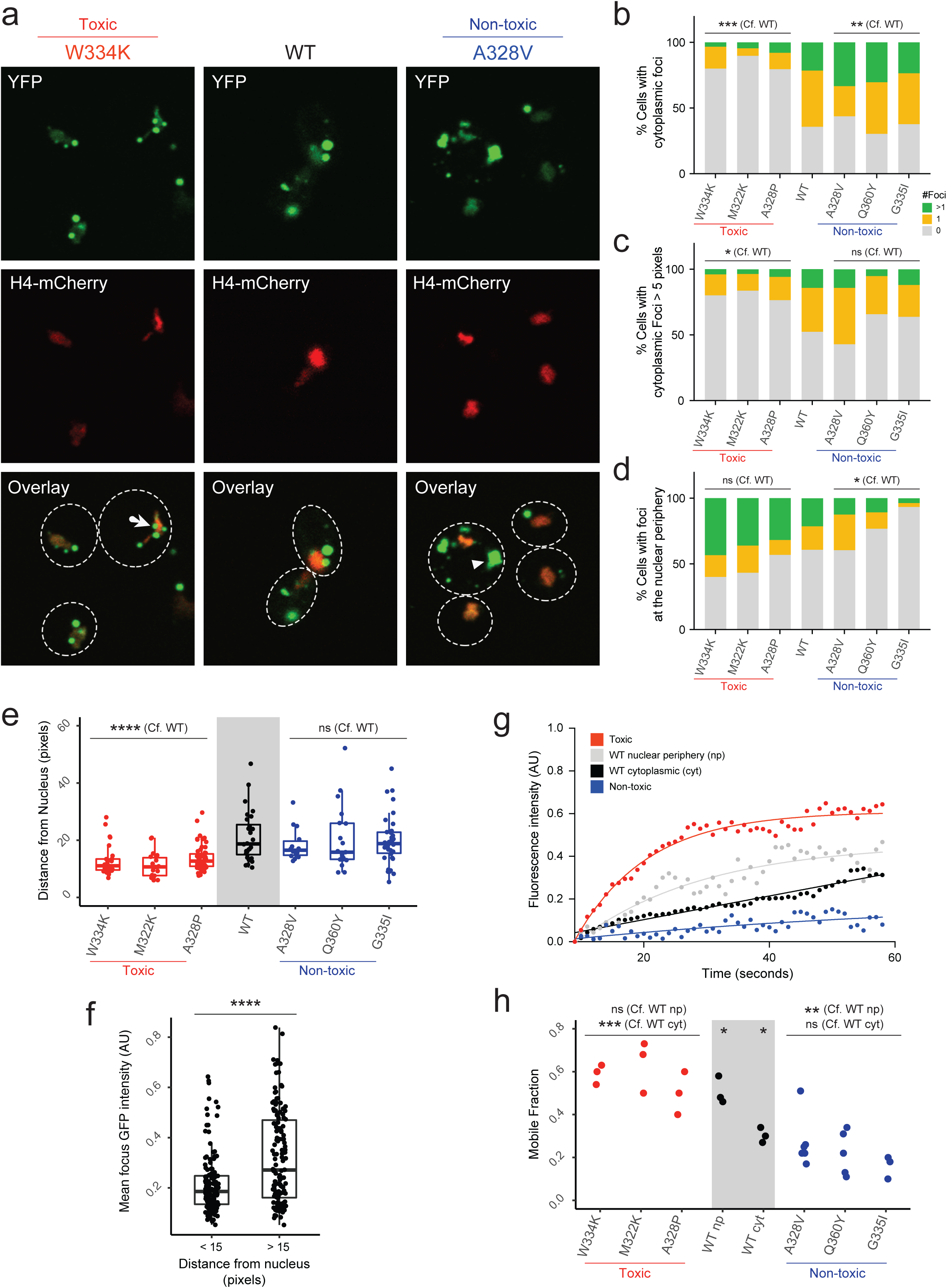
Mutations leading to formation of solid-like aggregates rescue toxicity. **a** Representative fluorescence microscopy Images of yeast cells expressing indicated YFP-tagged TDP-43 variants (W334K TDP-43 = toxic, A328V TDP-43 = non-toxic). H4-mCherry marks nuclei (red). Contrast was enhanced equally for the green and red channels in all images. **b** Percentage of cells with cytoplasmic foci (Cells scored: n[toxic]=219, n[WT]=30, n[non-toxic]=213). Fisher’s Exact test. **c** Percentage of cells with cytoplasmic foci with size over 5 pixels automatically detected by CellProfiler. Fisher’s Exact test. (Cells scored: n[toxic]=167, n[WT]=23, n[non-toxic]=167) **d** Percentage of cells with foci at the nuclear periphery (Cells scored: n[toxic]=219, n[WT]=30, n[non-toxic]=213). Fisher’s exact test. **e** Distance of foci from nucleus center for toxic (red), non-toxic (blue), and WT (black) TDP-43. Boxplots represent median values, interquartile ranges and Tukey whiskers with individual data points superimposed. Kruskal Wallis with Dunn’s multiple comparisons test (n = >20 foci / variant). **f** Average fluorescence intensity of foci localised closer (<15 pixels, n=147) or further (> 15 pixels, n=138) from the nucleus. Boxplots represent median values, interquartile ranges and Tukey whiskers with individual data points superimposed. Mann-Whitney test. **g** Representative individual fluorescence recovery traces for variants reported in panel e. Lines are the result of a single exponential fitting. **h** Mobile Fraction as calculated by fitting FRAP traces for toxic (red), non-toxic (blue) and WT (black) TDP-43. Each point results from fitting an individual trace. One-way ANOVA with Tukey’s multiple comparisons test. Images were taken on cells growing from at least 3 independent starting colonies. * *P* < 0.05, ** *P* < 0.01, *** *P* < 0.001,**** *P* < 0.0001.

**Figure 4.**
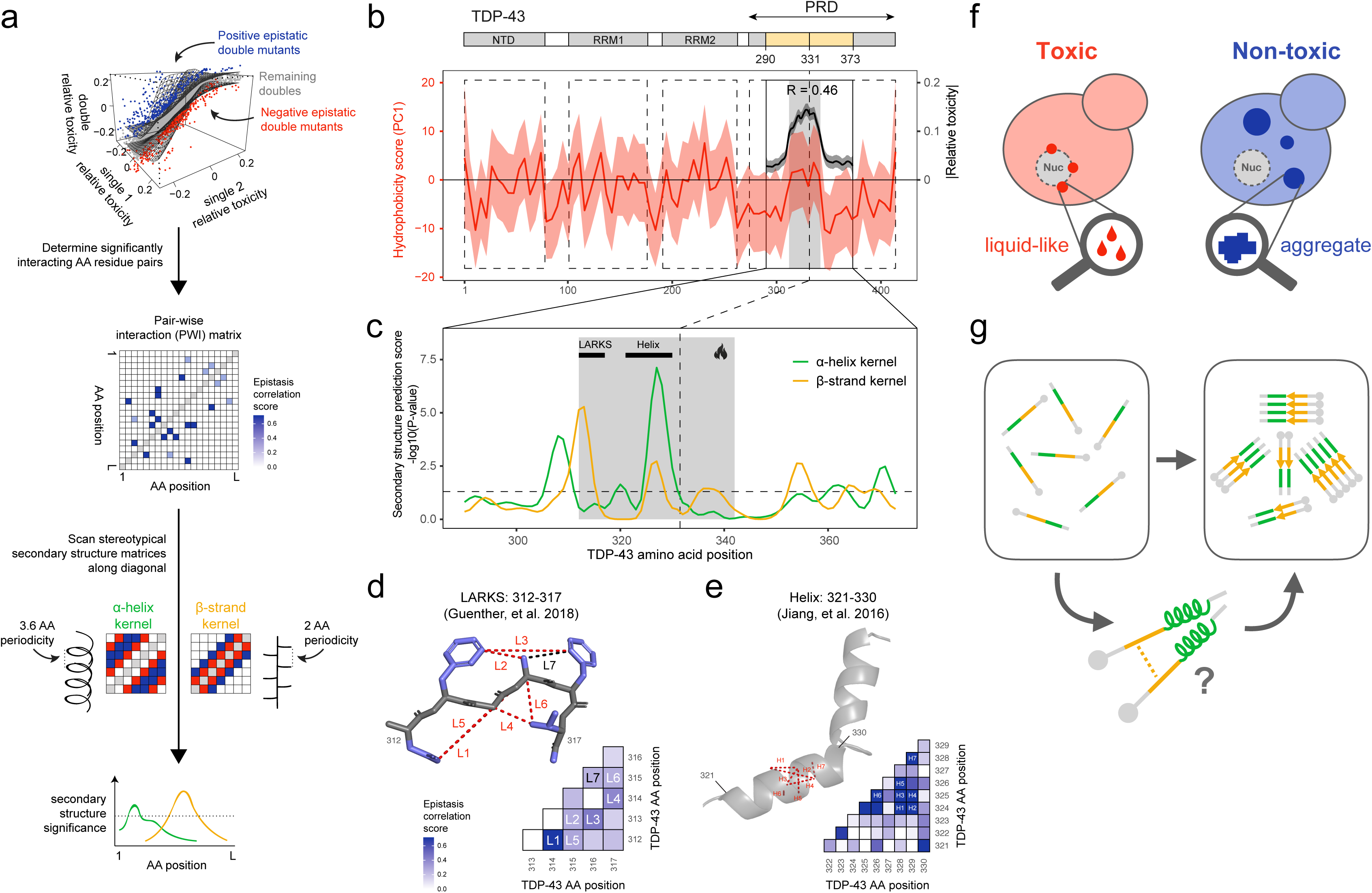
Correlated patterns of epistasis predict secondary structural elements within the PRD of TDP-43. **a** Schematic representation of the computational strategy to identify *in vivo* secondary structures. Double mutant variants are classified as epistatic if they are more (95^th^ percentile) or less (5^th^ percentile) toxic than other variants with similar single mutant toxicities (top). A pair-wise interaction (PWI) matrix of epistasis correlation scores is then constructed by quantifying the similarity of a pair of positions interactions with all other mutated positions in the protein. The epistasis correlation scores along the diagonal of the PWI matrix are then tested for agreement with the stereotypical periodicity of ⍰-helix and β-strand, using two-dimensional kernels (bottom), to calculate the likelihood of adjacent positions forming secondary structures. **b** Local polynomial regression (loess) of hydrophobicity (PC1) of the WT TDP-43 sequence with 95% confidence interval. For reference, smoothed toxicity estimates in the mutated positions within the PRD are shown. The Pearson correlation coefficient (R) between hydrophobicity and mean toxicity effects of single mutants at each position before smoothing is indicated. **c** Secondary structure predictions from epistasis correlation scores for ⍰-helix and β-strand kernels based on the strategy described in panel a. Black bars annotate previously described structural features: LARKS, low-complexity aromatic-rich kinked segment (312-317)^53^; Helix (321-330)^37^. The dashed horizontal line indicates the nominal significance threshold P=0.05. **d** Epistatic interactions in region 312-317 are consistent with positions of similar side-chain orientations interacting in a previously reported *in vitro* LARKS structure. Epistasis correlation matrix and top seven epistasis correlation score interactions annotated on the LARKS reference structure (monomer from PDB entry 5whn). Dashed lines on structure connect interacting residues at minimal distance between side chain heavy atoms. Side chain atoms are depicted in blue. **e** Epistatic interactions in region 321-330 are consistent with positions of similar side-chain orientations interacting in an ⍰-helix. Epistasis correlation matrix and top seven epistasis correlation score interactions annotated on the helix reference structure (monomer from PDB entry 5whn). **f** Model of how AA changes determine toxicity of TDP-43: mutations that promote formation of insoluble cytoplasmic aggregates decrease its toxicity, while mutations that cause the protein to stall in a liquid de-mixed phase increase TDP-43 toxicity to the cell **g** Secondary structure elements, within the toxicity hotspot 312-342, promote the aggregation process of TDP-43, with a transient helix forming on pathway to β-rich aggregates.

That increased hydrophobicity and aggregation potential are strongly associated with reduced toxicity across >50,000 genotypes was unexpected given previous work that reported an increased number of intracellular aggregates for a set of TDP-43 variants toxic to yeast^55^ and the widely-held view that aggregation is harmful to cells^44,53,57^. We therefore further investigated the effects of mutants that alter the hydrophobicity and toxicity of TDP-43.

WT TDP-43 localizes to both the nucleus and to the cytoplasm of yeast cells^55,56^ (Fig. 3a). In the nucleus, TDP-43 is diffuse, but in the cytoplasm it forms *puncta*, consistent with previous observations^43,58^. We observed that cytoplasmic WT TDP-43 forms two types of assemblies: small foci in the nuclear periphery and larger foci detached from the nucleus (Fig. 3a-c). We found that mutations that decrease TDP-43 hydrophobicity and increase TDP-43 toxicity increase the number of the small foci at the nuclear periphery and reduce the number of large distal foci (Fig. 3b, c, f, ED Fig. 5a). In contrast, mutations that increase hydrophobicity and reduce toxicity reduce the number of small nucleus-associated foci and increase the number of large distal foci (Fig. 3b, c, f, ED Fig. 5a).

We used fluorescence recovery after photobleaching (FRAP) to characterize the dynamics of TDP-43 variants in the different foci. The large cytoplasmic foci formed by non-toxic variants show little exchange of TDP-43 molecules with the soluble cytoplasmic pool. In contrast, the small foci localized at the nuclear periphery can exchange more protein with the cytoplasm, consistent with a more liquid-like state (Fig. 3d, e). Such differences in dynamics have been previously described for distinct types of misfolded protein compartments^59^. The large immobile TDP-43 foci are also brighter than the small dynamic ones (Fig. 3g), similar to what has been observed for Huntingtin variants that partition between immobile bright assemblies and liquid-like dimmer ones^60^. The non-toxic TDP-43 variants also had a higher protein concentration quantified by Western blotting (ED Fig. 5b).

Taken together, these results suggest that mutations that increase the hydrophobicity of TDP-43 result in a re-localization of the protein away from small and dynamic, liquid-like foci at the nuclear periphery to large and more solid aggregates in the cytoplasm. A reduction in hydrophobicity has the opposite effect.

The hotspot region of the TDP-43 PRD (AA 312-342) contains a conserved region^37,38^, with hydrophobicity more similar to the globular domains of TDP-43 than to the surrounding hydrophilic disordered regions (Fig. 4b). The hotspot is contained within a region (311-360) that was previously shown to be sufficient for both *in vitro* aggregation and the formation of cytoplasmic foci^37^. Fragments from within this region have previously been shown to have the potential to form different types of secondary structures *in vitro*. More specifically, nuclear magnetic resonance (NMR) spectroscopy of the PRD revealed that residues 321-342 can adopt an ⍰-helical structure in certain conditions^37,38,49^ and four different 6-11 AA peptides from the region could form cross-β amyloid or amyloid-like fibrils whose structures were determined by X-ray crystallography^53^. However, it is unknown whether any of these structures exist *in vivo* for full length TDP-43.

We have shown recently that the pattern of genetic (epistatic) interactions between mutations in a protein can determine the structure of that molecule when it is performing the function that is being selected for^52^. In particular, when a sequence forms an ⍰-helix, the side chains of residues separated by 3-4 AA are close in space and similarly oriented so that mutations in these AA interact similarly with mutations in the rest of the protein. In contrast, in a β-strand, the side chains of residues separated by 2 AA are close and similarly oriented and so make similar genetic interactions with other mutations (Fig. 4a).

We used 52,272 double mutants (excluding STOP codon variants) in our dataset to identify pairs of mutations that genetically interact. We first identified pairs of mutations that had unexpectedly high or low toxicity (<5^th^ and >95^th^ percentile of the expected toxicity distribution, negative and positive epistasis for growth rate, respectively). We then quantified the similarity of epistasis enrichment profiles between pairs of positions and compared these patterns to those expected for ⍰-helices and β-strands, scoring significance by randomisation^52^ (Fig. 4a).

This revealed that the patterns of epistasis in our dataset are consistent with two secondary structure elements forming inside the PRD *in vivo*: a β-strand at residues 311-316 and an ⍰-helix at residues 324-331 (Fig. 4c). The β-strand identified by the epistasis analysis coincides with one of the peptides in the TDP-43 PRD that, *in vitro*, can form cross-β structures^53^ typical of protein aggregates (Fig. 4d). The crystals of this specific peptide consist of a non-conventional β-strand termed a low-complexity aromatic-rich kinked segment (LARKS)^61^. In this *in vitro* structure, Phe 313 and Phe 316 face the same side of the sheet, whereas in a canonical sheet the side chains of odd and even residues face opposite sides. Strikingly, this non-canonical contact between Phe 313 and Phe 316 is also identified by the *in vivo* epistasis analysis, with a similarity in interaction profile ranking amongst the top two residue pairs in this region. In addition, the contact between Phe 316 and Ala 315, which again is compatible with a LARKS but not with a canonical β-strand, has the highest predicted contact score among neighboring residues (Fig. 4d). The predicted contact map built on the basis of *in vivo* epistatic interactions strikingly matches the PDB structure for LARKS 312-317 (Fig. 4d, ED Fig. 6).

On the other hand, the genetic interactions of mutations in the 324-330 region match those expected for an ⍰-helix (Fig. 4e). This region is part of the portion (321-342) of TDP-43 that can transiently and cooperatively fold into a ⍰-helix *in vitro*^*38,49,62*^. This helix is stabilized by inter-molecular contacts and its self-interaction was proposed to seed liquid-demixing *in vitro*. Amyloid fibrils can grow from the liquid de-mixed state and circular dichroism spectroscopy revealed that the helix transitions to a β-sheet over time, compatible with the process of aggregation^37,62^. On the basis of epistasis, the top scoring predicted contacts in this region are between residues separated by 3-4 AA such as Ala 324 and Ala 328, or Ala 325 and Ala 328, consistent with interactions between side chains of an ⍰-helix (Fig. 4e).

The pattern of *in vivo* epistatic interactions between mutations in TDP-43 therefore is compatible with a model in which two of the secondary structures that have previously been observed *in vitro* for fragments of TDP-43 actually form *in vivo* in the full-length protein. This indicates that the ‘unstructured’ PRD of TDP-43 is at least partially structured *in vivo*. A parsimonious model based on previous *in vitro* work^37^ is that the helix forms first in the pathway of aggregation towards a β-rich species (Fig. 4g). Consistent with this, destabilizing mutations, such as any substitution of Phe 313 and Phe 316 in the LARKS, or the introduction of proline into the 324-330 helix, increase toxicity (Fig. 1f).

Taken together, by quantifying the effects of >50,000 mutants of TDP-43, we have found that aggregation of TDP-43 is actually protective to yeast cells. We propose that this is because aggregation titrates TDP-43 away from a toxic liquid-like phase at the nuclear periphery (Fig. 4f). Liquid de-mixed TDP-43 was recently shown to recruit the nuclear pore component Nup62 and the importin-⍰ transporter, resulting in nuclear transport impairment^46^. That TDP-43 aggregates are protective rather than toxic is consistent with previous work, including the rescue of toxicity by the accumulation of RNA lariats that sequester TDP-43 into large aggregates^63^.

Mutations in TPD-43 had their strongest effects on toxicity within a hydrophobic region where the patterns of genetic interactions are consistent with the *in vivo* formation of two structural elements that have been shown to be important for the phase separation and aggregation of fragments of TDP-43 *in vitro*^38,49, 53, 62^.

More generally, our results show that deep mutagenesis is a powerful approach for determining the sequence-function relationships of intrinsically disordered proteins, including their *in vivo* structural conformations. The conformations of ‘unstructured’ proteins are notoriously difficult to study and the interactions between mutations in double mutants provide a general method to probe the *in vivo* structures of these proteins whenever a selection assay is available. We envisage that this approach can be adopted to study the functions, toxicity, and *in vivo* structures of other intrinsically disordered proteins, including the many other proteins implicated in neurodegenerative diseases.

Our conclusions derived from deep mutagenesis of TDP-43 are also consistent with observations for other ALS genes, including the reduced toxicity of SOD-1 variants that increase aggregation^11,64^. They are also consistent with the increasing evidence that insoluble aggregates are not pathogenic in multiple other neurodegenerative diseases^4,65,66^, and with the clinical failure of therapeutic approaches that reduce aggregation *in vitro* and *in vivo*^12,13,19,67,68^. Indeed, if insoluble aggregates titrate proteins away from alternative toxic phases, then promoting rather than alleviating aggregation might be the appropriate therapeutic goal.

## Methods

### Yeast Strains and Plasmids

*Saccharomyces cerevisiae* S288C BY4741 (*MATa his3*Δ*1 leu2*Δ*0 met15*Δ*0 ura3*Δ*0*) was used in all experiments. Plasmid pRS416 containing TDP-43 or TDP-43-YFP under control of the Gal1 promoter was purchased from Addgene^55^. Mutagenesis for the characterization of TDP-43 variants was performed through PCR linearization with specifically-designed primers (Supplementary Table 1, primers: BB_1 to BB_6). The resulting products were then either treated with DpnI or purified from a 1% agarose gel with a QIAquick Gel Extraction Kit (Qiagen) and transformed into *E.*coli DH5α competent cells (Invitrogen) for plasmid purification and validation through Sanger sequencing.

### Library construction

Two 186 nt oligonucleotides were purchased from TriLink. Each consisted of a ‘doped’ region of 126 nt, corresponding to TDP-43 AA 290-331 or AA 332-373, flanked by 30 nt of the WT TDP-43 sequence on each side. Each position in the mutated area, was doped with an error rate of 1.59%. The target frequency for each library was 27.0% for single mutants and 27.3% for double mutants. With this approach, the WT sequence was represented with a frequency of 13.3%. Each oligonucleotide was amplified by PCR (Q5 High-Fidelity DNA Polymerase, NEB) for 15 cycles, purified using an E-gel electrophoresis system (Agarose 2%) followed by column purification with a MinElute PCR Purification Kit (Qiagen). In order to introduce the doped sequence in the full-length TDP-43 sequence the purified oligonucleotide was cloned into 100 ug of linearized pRS416 Gal TDP-43 by a Gibson approach (Supplementary Table 1, primers BB_7 to BB_14). The product was then transformed into 10-beta Electrocompetent *E. coli* (NEB), by electroporation in a Bio-Rad GenePulser machine (2.0k V, 200 Ω, 25 μF). Cells were recovered in SOC medium (NEB) for 30 min and plated on LB with ampicillin. A total of ∼2.7×10^6^ transformants were estimated. The plasmid library was purified with a GeneJET Plasmid Midiprep Kit (Thermo Scientific).

### Yeast Transformation and Selection experiments

Yeast cells were transformed with the TDP-43 doped plasmid in 4 independent biological replicates for each library. One single colony was grown overnight in 30 ml YPDA medium at 30°C for each replica. Cells were diluted to 0.3 OD 600 nm in 175 ml of YPDA and incubated for 4 h at 30°C. Cells were then harvested, washed, re-suspended in 8.575 mL SORB (100 mM LiOAc, 10 mM Tris pH 8.0, 1 mM EDTA, 1 M sorbitol) and incubated for 30 min at room temperature. For the transformation, 10 mg/mL of salmon sperm DNA and 3.5 ⍰g TDP-43 plasmid library were used. Cells were mixed to 100 mM LiOAc, 10 mM Tris-HCl pH 8.0, 1 mM EDTA/NaOH pH 8.0 and 40% PEG 3350. Heat-shock was performed for 20 min at 42°C. YPD with 0.5 M sorbitol was used to recover the cells, incubating them for 1 h at 30°C. After recovery, cells were resuspended in SC-URA 2% raffinose medium, while an aliquote was plated to calculate transformation efficiency.

After ∼50 h of growth, cells were diluted in SC-URA 2% raffinose medium and grown for 4.5 generations. At this stage, 400 mL of each replica were harvested, washed, split into two tubes and frozen at −20°C for later extraction of input DNA. To induce plasmid expression, for each replicate two cultures were diluted in SC-URA 2% galactose medium. After 5-6 generations, 2×400 mL for each replicate were harvested to obtain output pellets for DNA extraction.

### DNA Extraction and Library Preparation

Input and Output pellets were resuspended in 1.5 mL extraction buffer (2% Triton-X, 1% SDS, 100 mM NaCl, 10 mM Tris-HCl pH 8.0, 1 mM EDTA pH 8.0). Two cycles of freezing in an ethanol-ice bath and heating at 62°C were performed. Deproteinization was performed using 25:24:1 phenol-chloroform-isoamyl alcohol and glass beads. After centrifugation, the aqueous phase, containing the DNA, was recovered and treated again with phenol-chloroform-isoamyl alcohol. The samples were incubated 30 min at −20°C with 1:10V 3M NaOAc and 2.2V 100% ethanol for DNA precipitation. At this stage and after centrifugation for 30 min, the pellets were dried overnight at room temperature. RNA was eliminated by incubation with RNAse 10 mg/mL for 30 min at 37°C. DNA purification was achieved with a QIAEX II Gel Extraction Kit (Qiagen) and DNA was eluted in 375 μL of elution buffer. DNA concentration was measured by q-PCR, with primers annealing to the Ori site of the pRS416 plasmid (Supplementary Table 1, primers BB_15, BB_16).

The TDP-43 library was then prepared for deep sequencing by PCR amplification in two steps using Q5 High-Fidelity DNA Polymerase (NEB). In step 1, 300 million plasmids were amplified for 15 cycles using frame-shifted adaptor primers with a partial homology to standard Ilumina sequencing primers (Supplementary Table 1, primers BB_17 to BB_51). Samples were treated with ExoSAP (Affymetrix) and purified with QIAEX II kit (Qiagen). PCR products from the first step were used as templates in the second PCR step, where indexed Illumina primers (Supplementary Table 1, primers TS_HT_D7X_7 to TS_HT_D7X_95) were used for a 10 cycles amplification. DNA concentration was then quantified by means of a Quant-iT™ PicoGreen® dsDNA Assay Kit (Promega). All replicates were pooled together in an equimolar ratio. Finally, the pooled sequencing library was run on a 2% agarose gel, purified and sent for 125 bp paired-end Illumina sequencing at the CRG Genomics Unit.

### Individual Growth Rate Measurements

Yeast cells expressing selected TDP-43 variants were grown overnight in SC-URA 2% raffinose non-inducing medium and diluted to 0.2 OD 600 nm until exponential phase. Then they were diluted to 0.1 OD 600 nm in SC-URA 2% galactose to asses growth in inducing conditions. Growth was monitored by measuring OD 600 nm in a 96-well plate at 10 min intervals inside an Infinite M200 PRO microplate reader (Tecan). Plates were kept constantly shaking at 30°C. Growth curves were fitted in order to extrapolate growth rates that correspond to the maximum sloped of the linear range of the LN(OD) curve over time.

### Equipment and Settings

Imaging was performed by using a Confocal TCS SP8 and a Confocal TCS SP5 (Leica) equipped with PMT detectors both for fluorescence and transmitted light images. AOBS beam-splitter systems are in place on both instruments. 63X oil immersion objectives and the LAS AF software were used for all imaging. YFP fluorescence was excited with a 488 nm laser, while mCherry fluorescence with a 561 nm laser. Ranges for emission detection were 495-554 and 637-670 nm respectively. Image depth is 8-bit in all cases and pixel size equals 120.4 nm. The LUT is linear and covers the full range of the data.

### Fluorescence Microscopy and Image Analysis

Yeast cells expressing TDP-43 selected variants were grown in SC-URA 2% raffinose non-inducing medium and then transferred to SC-URA 2% galactose medium to induce protein expression for 8h. They were then imaged under a Confocal TCS SP8 microscope (Leica). Counting of foci was conducted both manually and by automated pipelines using the CellProfiler software where quantification of fluorescent intensity was tracked for each focus. The coordinates of the center of each focus and nucleus were also derived from CellProfiler and used to calculate distances using a custom R script (pipelines available at https://github.com/lehner-lab/tardbpdms_cellprofiler_scripts).

### Fluorescence Recovery after Photobleaching (FRAP)

Yeast cells expressing TDP-43 selected variants were grown in SC-URA 2% raffinose non-inducing medium and then transferred to SC-URA 2% galactose medium to induce protein expression for 8h. The cells were immobilized to an 8-well cover slide by Concanavalin-A-mediated cell adhesion. Cells were then imaged under a Confocal TCS SP5 microscope (Leica) where bleaching was achieved with 488 Laser Power at 70% for three frames (1.3 s/frame) while fluorescence recovery was recorded for 50 frames. The curves were then fitted to a single exponential, following normalization, with the EasyFrap package^69^.

### Protein Extraction and Western Blotting

Single yeast colonies were grown overnight in non-inducing medium and then diluted to 0.2 OD 600 nm in Galactose medium to induce protein expression for ∼8 h. At this stage, 6×10^7^ cells were collected and re-suspended in 200 μL EtOH and 2.5 μL PMSF. Samples were vortexed with glass beads for 15 min at 4°C and frozen overnight at - 80°C. The samples were dried in a speed vacuum for 20 min and resuspended in 200 μL solubilizing buffer (20mM Tris HCl pH 6.8,2% SDS). After boiling for 5 min, the lysate fraction was run on a NuPAGE 4-12% Bis-Tris gels (Novex) and transferred to PVDF membranes in an iBlot (Invitrogen). Membranes were blocked with 5% milk powder in TBS-T and incubated overnight at 4°C with primary antibodies: anti-GFP mouse antibody (Santa Cruz sc-9996) and anti-PGKD1 mouse antibody (Novex 459250) diluted 1:1,000 and 1:5,000 in 2.5% powder milk respectively. Secondary antibody anti-proteinG was incubated for 1h at room temperature. Proteins were detected with an enhanced chemi-luminescence system (Millipore Luminata) and visualized using an Amersham Imager 600 (GE Healthcare).

### Sequencing data pre-processing

FastQ files from paired-end sequencing of replicate deep mutational scanning (DMS) libraries before (‘input’) and after selection (‘output’) were processed using a custom pipeline (https://github.com/lehner-lab/DiMSum, manuscript in prep.). DiMSum is an R package that wraps common biological sequence processing tools including FastQC (http://www.bioinformatics.babraham.ac.uk/projects/fastqc/) (for quality assessment), cutadapt (for demultiplexing and constant region trimming), USEARCH^70^ (for paired-end read alignment) and the FASTX-Toolkit (http://hannonlab.cshl.ed/fastx_toolkit/). First, 5’ constant regions were trimmed, but read pairs were discarded if 5’ constant regions contained more than 20% mismatches to the reference sequence. Read pairs were aligned (reads that did not match the expected 126bp length were discarded) and Phred base quality scores of aligned positions were calculated using USEARCH. Reads that contained base calls with Phred scores below 30 (290-331 DMS library) or below 25 (332-373 DMS library) were discarded. Approximately five and seven million reads passed these filtering criteria in each sample corresponding to the 290-331 and 332-373 libraries respectively. Finally, unique variants were counted and merged into a single table of variant counts (aggregated across technical output replicates) per DMS library. One out of four input replicates (and all associated output samples) from each DMS library were discarded due to considerably lower correlations with the other replicates (ED Fig. 1a, b).

### Variant toxicity and error estimates

All analyses of toxicity were performed on variants with a maximum of two AA mutations, but no synonymous mutations in other codons. Firstly, sample-wise counts for variants identical at the AA level were aggregated. For each replicate selection, relative toxicity of variants was calculated from variant counts in input 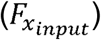 and output 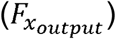 samples as 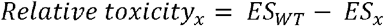, where 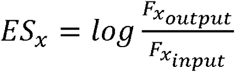 and *ES*_*WT*_ represents the WT enrichment score. Uncertainty of toxicity values was estimated as a combination of expected Poisson error based on read counts and error between replicate selections as 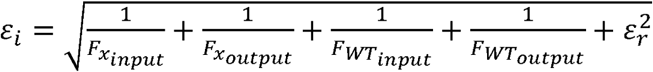. Here, *ε*_*r*_, the error between replicate selections, is estimated from the variance of toxicity estimates across replicates for variants whose expected count-based Poisson error approaches zero. Toxicity estimates and associated errors per replicate selection were also normalized by the replicate-specific number of cell doublings during selection to yield relative growth rates per generation.

In ‘doped’ variant libraries, individual double mutants are represented less frequently than single mutants or the WT sequence and due to this under-representation toxic double mutants (that are depleted due to slower growth during selection) are often not observed in the output samples (ED Fig. 1c). To calculate toxicity estimates for such double mutants and avoid skewed marginal toxicity distributions due to these drop-out events, we used a Bayesian approach to estimate toxicity of double mutants based on a prior, i.e. toxicity distributions of highly represented doubles that originate from single mutants with similar toxicity estimates^52^. These corrected toxicity estimates show improved heteroscedasticity and reduced variance, especially for under-represented double mutants (ED Fig. 1c).

Variant toxicity distributions were first normalized between replicate selections of the same DMS library to have equal standard deviations. Then toxicity estimates of each variant across replicate selections were merged by taking the error-weighted mean across replicate selections. Finally, distributions of merged toxicity estimates from each DMS library were centered on the error-weighted means of toxicity of single codon synonymous (silent) variants in each DMS library and scaled such that the error-weighted means of single STOP codon variants coincided for both DMS libraries (ED Fig. 2a-c). Furthermore, we removed low confidence variants supported by an average of less than ten input reads from all downstream analyses.

### Simple linear regression models to predict variant toxicity

We used simple linear regression to predict variant toxicity from (i) a collection of AA property features, (ii) a panel of scores from aggregation/structure algorithms and (iii) location with respect to the toxicity hotspot.

The AA property features were derived from a principal component analysis (PCA) of a curated collection of numerical indices representing various physicochemical and biochemical properties of AAs (http://www.genome.jp/aaindex/). From a total of 539 indices, we retained 379 high confidence indices with no missing values (including 5 additional indices absent from the original database; see Supplementary Table 2). Results of PCA and selected variable loadings on the normalized matrix are shown in ED Fig. 3. For single mutant variants, AA property feature values represent the difference between the WT and mutant PC scores.

Similarly, aggregation, disorder, structure and other feature values for single mutant variants represent the difference between scores obtained using WT and single mutant AA sequences. AGADIR, *cat*GRANULE and Tango provide a single score per AA sequence. Unless a single score per AA sequence was provided (i.e. AGADIR, *cat*GRANULE, Tango), individual residue-level scores were summed to obtain a score per AA sequence (i.e. BetaTPred3, DISEMBL, IUPred2A, Waltz, ZipperDB, Zyggregator). The entire PRD AA sequence was supplied to AGADIR and all unique six-mers to ZipperDB. For the remainder, the full-length AA sequence was used.

Variants inside the hotspot were defined as those with mutant residue positions in the range 312-342. Change in absolute charge (regardless of sign) is shown in Fig. 2d, e, because this feature is more predictive of toxicity than change in charge itself (not shown). For double mutant variants, we summed the feature values of the constituent singles for both AA property and aggregation/structure algorithm features. Regression models were built using either (i) all variants, restricting variants to those occurring either (ii) inside or (iii) outside the toxicity hotspot (for double mutants both mutations have to occur either inside or outside the hotspot region), or (iv) including a binary location variable (0: one/all outside, 1: one inside, one outside, 2: one/all inside toxicity hotspot) and a third term indicating the interaction between location and the AA property or aggregation/structure algorithm feature.

### Predicting secondary structure from epistasis

Epistasis is the non-independence of mutation effects, i.e. the toxicity of double mutants is different from that expected given the toxicity of their constituent single mutant variants. We have previously shown that epistasis between double mutants can result from structural interactions within proteins and therefore can be used to infer secondary and tertiary structural features^52^. In brief, double mutants were classified as epistatic if they had more extreme toxicity values (below 5^th^ percentile or above 95^th^ percentile) than other double mutants with similar single mutant toxicities, which was estimated from non-parametric surface fits of double mutant toxicity as a function of a two-dimensional single mutant toxicity space (Fig. 4a).

Double mutants close to the lower or upper measurement range limits (where the power to detect significant epistasis is reduced) were excluded from epistasis quantification. We calculated position-pair enrichments for epistatic double mutants resulting in a pair-wise enrichment matrix. Diagonal entries on this matrix were imputed as column-wise mean enrichments. An epistasis correlation score matrix was then derived from this enrichment matrix by calculating the partial correlation of epistasis interaction profiles (columns of the enrichment matrix) between all pairs of positions. The rationale for the correlation score is that structurally close positions within a protein should have similar epistatic interactions with all other positions in the protein. Calculating partial correlations additionally removes transitive interactions and was found to be superior over epistasis enrichments in estimating secondary structures^52^.

Secondary structure propensities were calculated by testing for agreement of epistasis correlation score patterns with the stereotypical periodicities of an ⍰-helix and β-strand, using two-dimensional kernels at each position along the diagonal of the epistasis correlation score matrix^52^. Significance of secondary structure propensities was assessed by comparison to propensities derived from 10^4^ randomized epistasis correlation score matrices.

Similarly, LARKS structure propensities were calculated using PDB-structure derived contact matrices based on a minimal side-chain heavy atom distance of 4.5Å (ED Fig. 6) for both WT (PDB entry: 5whn) and mutant sequences (PDB entries: 5whp and 5wkb). Contact matrix values were normalised to have zero sum. Association score matrix values were normalised to have mean of zero and unit variance. Significance of LARKS structure propensities was assessed by comparison to propensities derived from 10^4^ randomized epistasis correlation score matrices, where randomization was restricted to within-LARKS interactions i.e. distances compatible with a six-mer.

## Figure Legends

**Extended Data Figure 1. Quality control and pre-processing of DMS datasets. a** Correlation of mutant variant counts between all input replicates in DMS library 290-331. The Pearson correlation coefficients (R) are indicated in the upper matrix triangle. Replicate selection 2 was removed from downstream analyses due to lower than average correlation with other replicates **b** Correlation of mutant variant counts between all input replicates in DMS library 332-373. Replicate selection 1 was removed from downstream analyses due to lower than average correlation with other replicates. **c** Comparison of relative toxicity of single and double AA mutants and mean input read count for each DMS library before and after Bayesian correction of double AA mutant toxicity estimates. The vertical dashed line indicates the minimum mean input read count threshold (10) for variants used in downstream analyses. **d** Correlation of toxicity estimates between all retained replicates for single and double amino acid (AA) mutants from library 290-331. The Pearson correlation coefficients (Corr) are indicated. **e** Similar to panel d except showing results corresponding to retained replicates from library 331-373.

**Extended Data Figure 2. Inter-library normalization of DMS datasets and toxicity of human disease mutations. a** Toxicity distributions of single codon synonymous (silent) variants (top), single and double mutants (middle) and single STOP codon variants, shown separately for each library (see color key) and before centering and scaling. **b** As in panel a, but after inter-library normalization by centering on the error-weighted means of toxicity of single codon synonymous (silent) variants. **c** As in panel b, but after additionally scaling such that the error-weighted means of single STOP codon variants coincided. Mean toxicity of variants with single STOP codon mutation is indicated by dashed vertical line. **d** Relative toxicities of classified human disease AA substitution variants (colored dots; see key) below the relative toxicity distribution of all single and double AA mutants assayed in this study. Disease variants observed in more than one patient are classified as recurrent. **e** Table of all human disease AA substitution mutations in TDP-43 used in panel d.

**Extended Data Figure 3. Principle components analysis of amino acid physicochemical properties. a** Results of PCA of a curated collection of numerical indices representing various physicochemical and biochemical properties of AAs (see Methods). Biplot matrix indicating variable loadings of the top 5 PCs. Colors indicate text matches to index descriptions (see color key). **b** Screeplot indicating percentage variance explained by all PCs.

**Extended Data Figure 4. Linear regression models to predict mutant variant toxicity. a** Percentage variance of residual relative toxicity (after controlling for hydrophobicity and location) explained by linear regression models predicting single and double mutant variant toxicity from changes in AA properties upon mutation (left) and using scores from aggregation/structure algorithms (right). Different regression models were built for different subsets of the data. Simple linear regression models for all variants (blue) or only variants inside (red) or outside (yellow) the hotspot region. And a regression model using all variants and including a binary location variable (inside/outside hotspot) as well as an interaction term between binary location variable and the indicated AA property feature (green). **b** Percentage variance of relative toxicity explained by linear regression models predicting single mutant variant toxicity from changes in AA properties upon mutation (left) and using scores from aggregation/structure algorithms (right). **c** Similar to panel b except showing results using double mutant variants.

**Extended Data Figure 5. Scoring of intracellular phenotypes and expression of toxic and non-toxic mutants. a** Scoring of intracellular phenotypes by automated foci counting. Percentage of cells with foci at the nuclear periphery automatically scored by CellProfiler. Fisher’s Exact test. **b** Immunohistochemistry of toxic and non-toxic mutants. Expression of different TDP-43 variants after 8h induction of protein expression in Galactose was measured by Western Blotting. Phosphoglycerate Kinase 1 was used as a Loading Control.

**Extended Data Figure 6. LARKS structure propensities. a** Contact matrix based on a minimal side-chain heavy atom distance of 4.5Å derived from WT LARKS PDB structure 5whn. **b** Contact matrix derived from mutant LARKS PDB structure 5whp. **c** Contact matrix derived from mutant LARKS PDB structure 5wkb. **d** LARKS structure propensities for PDB-structure derived contact matrices shown in panels a-c (see Methods). The dashed vertical line indicates the start position of the LARKS (TDP-43 AA residue 312).

**Extended Data Figure 7. Toxicity of variants as a function of hydrophobicity and aggregation propensity. a** Toxicity of variants with single (left) or double (right) mutations occurring outside (top), inside (bottom) or in both locations (1 outside/1 inside) w.r.t. the hotspot region (middle), as a function of hydrophobicity changes (PC1) induced by mutation. The Pearson correlations (R) before binning are indicated. **b** Toxicity of variants with single (left) or double (right) mutations occurring outside (top), inside (bottom) or in both locations (1 outside/1 inside) w.r.t. the hotspot region (middle), as a function of changes in aggregation propensity (Zyggregator). The Pearson correlations (R) before binning are indicated.

## Data Availability Statement

Raw sequencing data and the processed data table (Supplementary Table 3) have been deposited in NCBI’s Gene Expression Omnibus and are accessible through the GEO Series accession number GSE128165 (https://www.ncbi.nlm.nih.gov/geo/query/acc.cgi?acc=GSE128165). All software code and custom scripts are available on GitHub: https://github.com/lehner-lab/DiMSum for raw read processing, https://github.com/lehner-lab/tardbpdms for all downstream analyses and to produce all figures, and https://github.com/lehner-lab/tardbpdms_cellprofiler_scripts for CellProfiler pipelines.

## Acknowledgments and Funding

Work in B.L.’s lab was supported by a European Research Council (ERC) Consolidator grant (616434), the Spanish Ministry of Economy and Competitiveness (BFU2017-89488-P), the AXA Research Fund, the Bettencourt Schueller Foundation, and Agencia de Gestio d’Ajuts Universitaris i de Recerca (AGAUR, SGR-831) G.G.T. was supported by the European Research Council (RIBOMYLOME_309545) and the Spanish Ministry of Economy and Competitiveness (BFU2014-55054-P and BFU2017-86970-P). We acknowledge support from the Spanish Ministry of Economy and Competitiveness, ‘Centro de Excelencia Severo Ochoa 2013-2017’, the EMBL Partnership, and the CERCA Program/Generalitat de Catalunya. We thank Pablo Baeza Centurión, Xavier Salvatella, Alexandros Armaos and Benjamin Lang for discussion and assistance and the Eisenberg lab for help with the Zipper DB analysis.

## Author Contributions

B.B. and B.L. conceived the project and designed the experiments; B.B. and M.S. performed the experiments; A.J.F, B.B. and J.M.S. performed analyses of sequences; A.J.F. and J.M.S. analysed the genetic interactions and structures; GGT initiated, designed and carried out the original computational analysis of physicochemical properties; B.B., A.J.F. and B.L wrote the manuscript with input from all authors.

## Author Information

Reprints and permissions information is available at www.nature.com/reprints. The authors declare no competing financial interests. Readers are welcome to comment on the online version of the paper. Correspondence and requests for materials should be addressed to B.B. and B.L.

## Supplementary Information

(Supplementary Table 1, Supplementary Table 2, Supplementary Table 3) is available in the online version of the paper

