## Supplementary figures for "The mutational landscape of a prion-like domain"

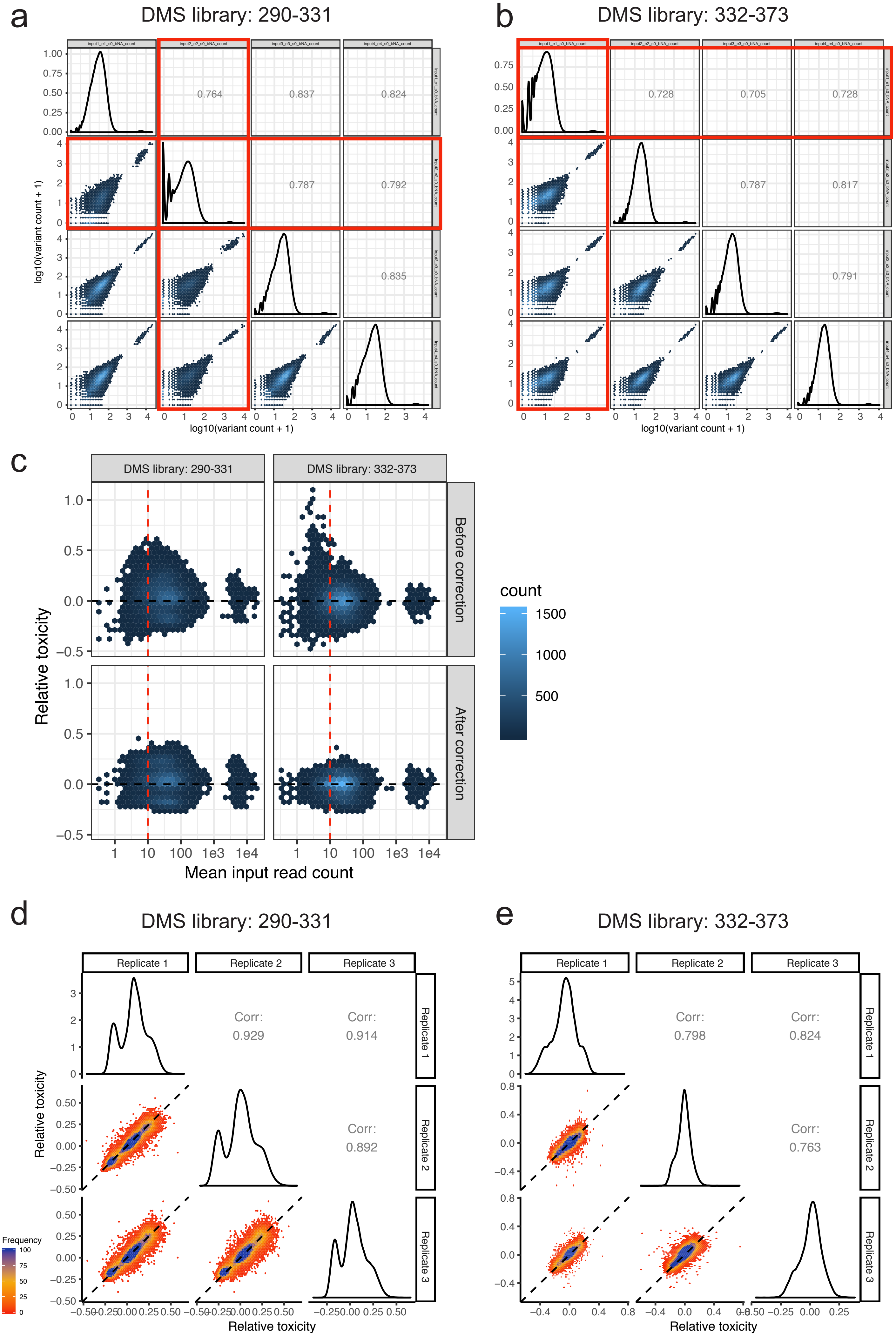


Extended Data Figure 1

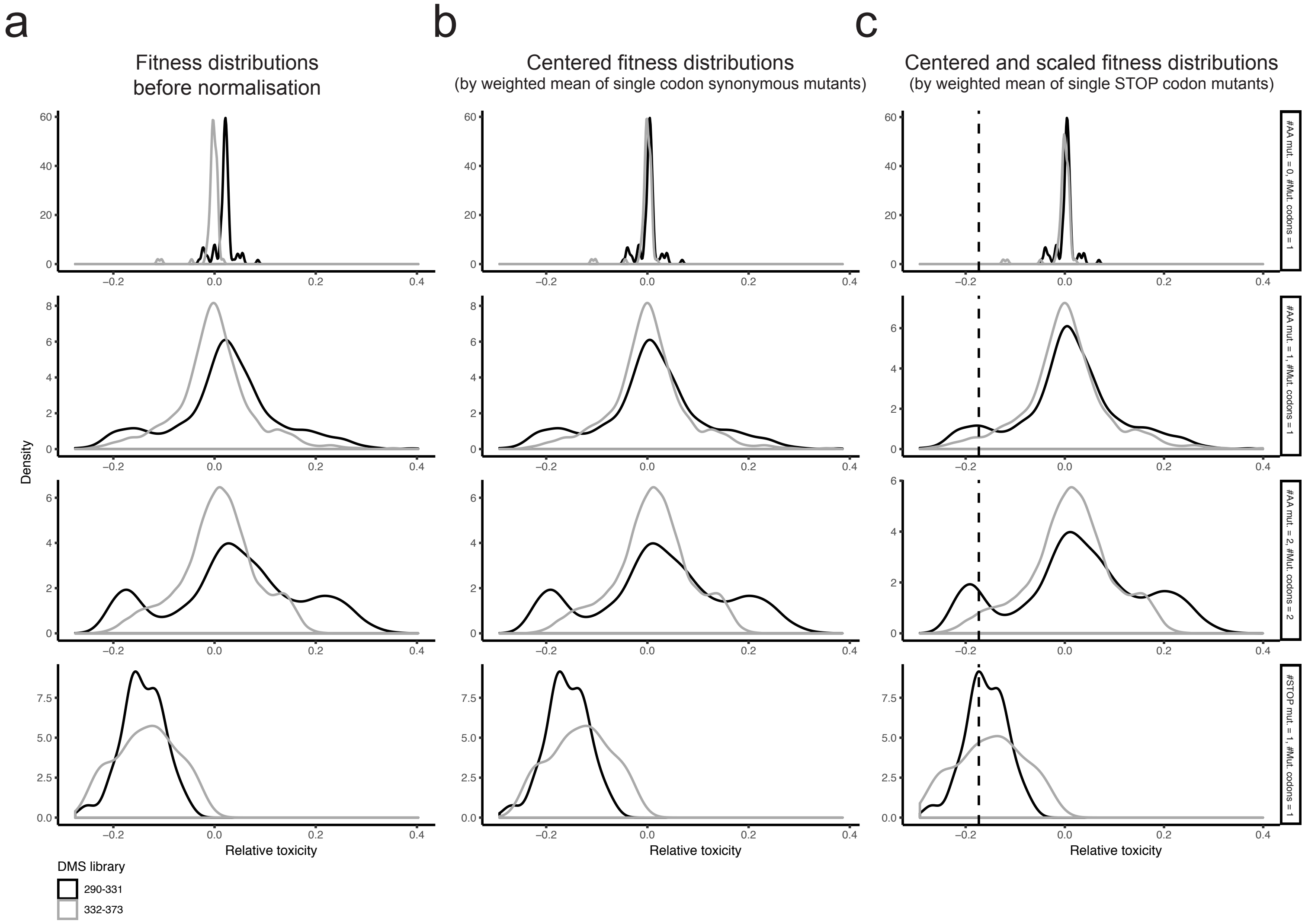


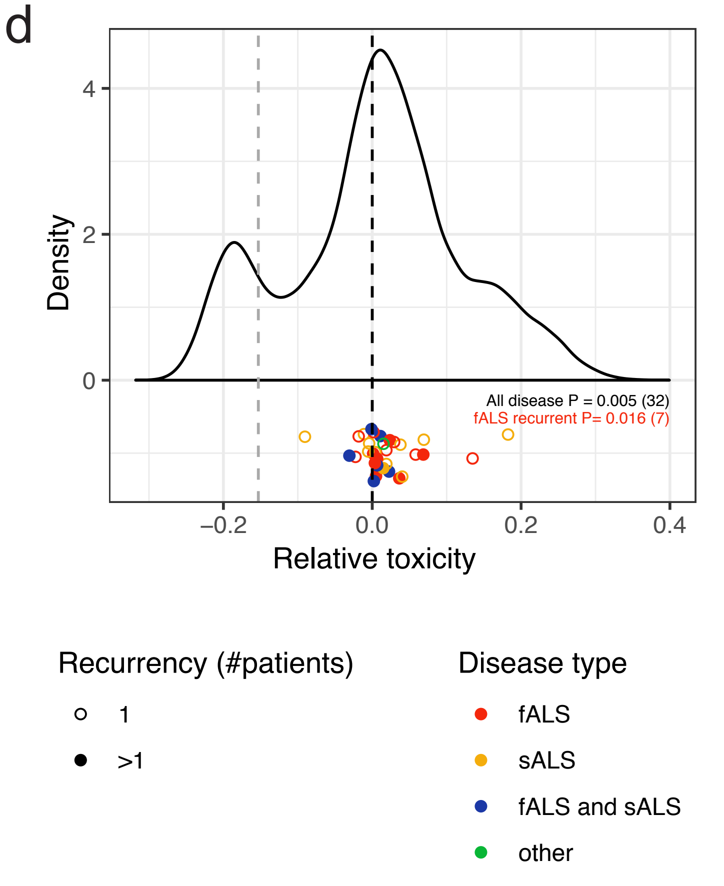


e

| Name | Position | #fALS | #sALS | Reference |
|---|---|---|---|---|
| Gly290Ala | 290 | 2 | | Van Deerlin (2008) Lancet Neurol 7, 409 |
| Ser292Asn | 292 | 1 | 2 | Xiong (2010) BMC Med Genet 11, 8 |
| Gly294Ala | 294 | | 1 | Sreedharan (2008) Science 319, 1668 |
| Gly294Val | 294 | 4 | 7 | Corrado (2009) Hum Mutat 30, 688 |
| Gly295Cys | 295 | 1 | | van Blitterswijk (2012) Hum Mol Genet 21, 3776 |
| Gly295Ser | 295 | 2 | 4 | Corrado (2009) Hum Mutat 30, 688 |
| Gly295Arg | 295 | | 1 | Corrado (2009) Hum Mutat 30, 688 |
| Gly298Ser | 298 | 2 | | Van Deerlin (2008) Lancet Neurol 7, 409 |
| Gln303His | 303 | | 1 | Lattante (2012) Neurology 79, 66 |
| Met311Val | 311 | 1 | | Lemmens (2009) J Neurol Neurosurg Psychiatry 80, 354 |
| Ala315Thr | 315 | 3 | | Gitcho (2008) Ann Neurol 63, 535 |
| Ala315Glu | 315 | 1 | | Fujita (2011) Neurology 77, 1427 |
| Ala321Val | 321 | | 1 | Kirby (2010) Neurogenetics 11, 217 |
| Ala321Gly | 321 | | 1 | Bäumer (2009) J Neurol Neurosurg Psychiatry 80, 1283 |
| Gln331Lys | 331 | | 1 | Sreedharan (2008) Science 319, 1668 |
| Ser332Asn | 332 | 1 | | Corrado (2009) Hum Mutat 30, 688 |
| Gly335Asp | 335 | | 1 | Corrado (2009) Hum Mutat 30, 688 |
| Met337Val | 337 | 7 | | Sreedharan (2008) Science 319, 1668 |
| Gln343Arg | 343 | 1 | | Rutherford (2008) PLoS Genet 4, e1000193 |
| Asn345Lys | 345 | 2 | | Rutherford (2008) PLoS Genet 4, e1000193 |
| Gly348Cys | 348 | 5 | 4 | Kabashi (2008) Nat Genet 40, 572 |
| Gly348Val | 348 | 2 | 2 | Kirby (2010) Neurogenetics 11, 217 |
| Gly348Arg | 348 | 1 | | Chiang, C. et al., Sci. Rep. 6, 21581, 2016 |
| Asn352Ser | 352 | 8 | 4 | Kühnlein (2008) Arch Neurol 65, 1185 |
| Asn352Thr | 352 | 2 | | Ticozzi (2011) Neurobiol Aging 32(11):2096-9 |
| Gly357Ser | 357 | | 2 | Iida (2012) Neurobiol Aging 33, 786 |
| Gly357Arg | 357 | 1 | | Chiang (2012) J Hum Genet 57, 316 |
| Met359Val | 359 | | | Borroni (2010) Rejuvenation Res 13(5):509-17 |
| Arg361Ser | 361 | | 1 | Kabashi (2008) Nat Genet 40, 572 |
| Arg361Thr | 361 | 2 | | Chiang (2012) J Hum Genet 57, 316 |
| Pro363Ala | 363 | | 1 | Daoud (2009) J Med Genet 46, 112 |
| Gly368Ser | 368 | | 1 | De Marco (2011) Acta Neuropathol 121, 611 |

### Extended Data Figure 2

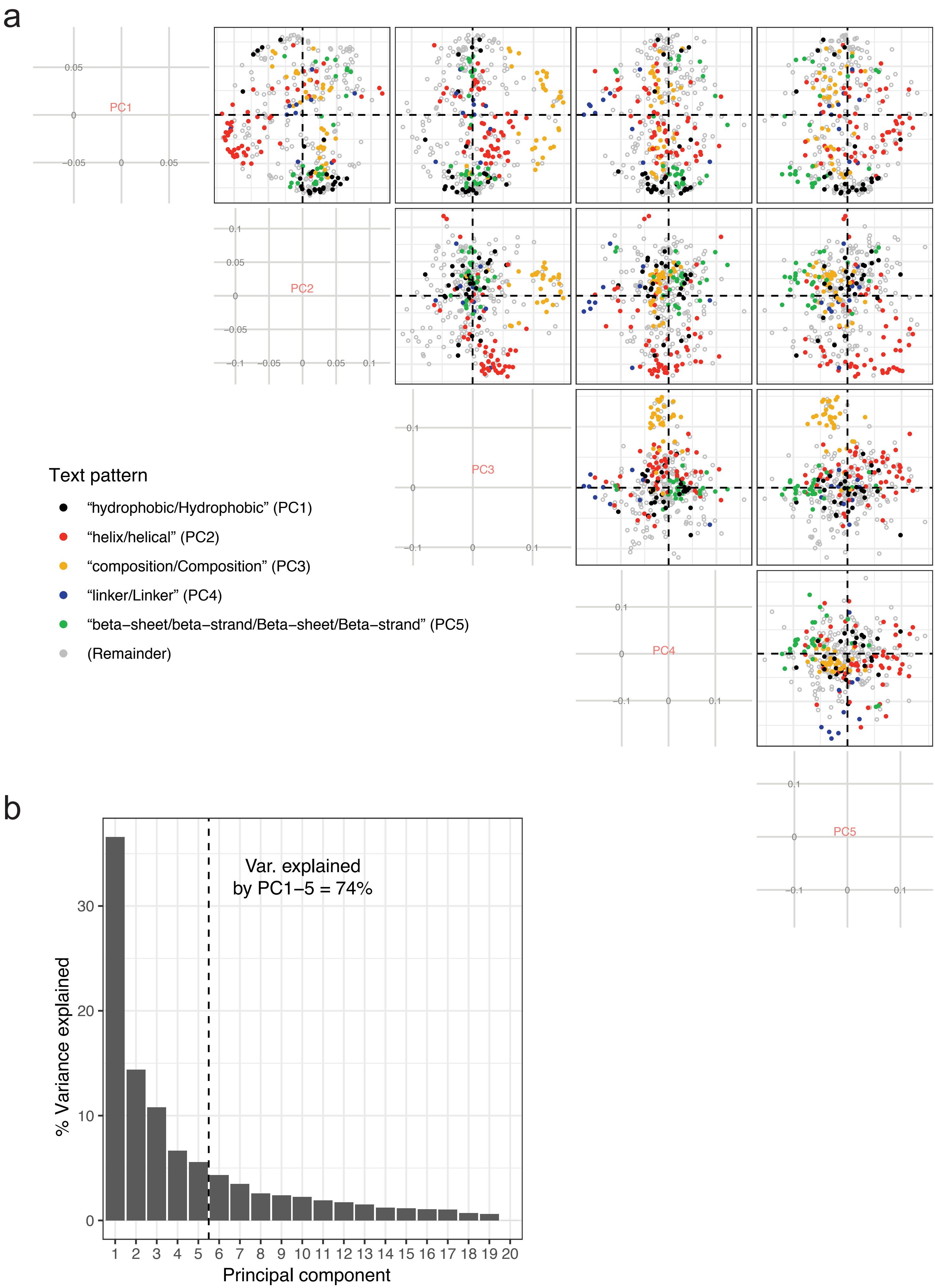


Extended Data Figure 3

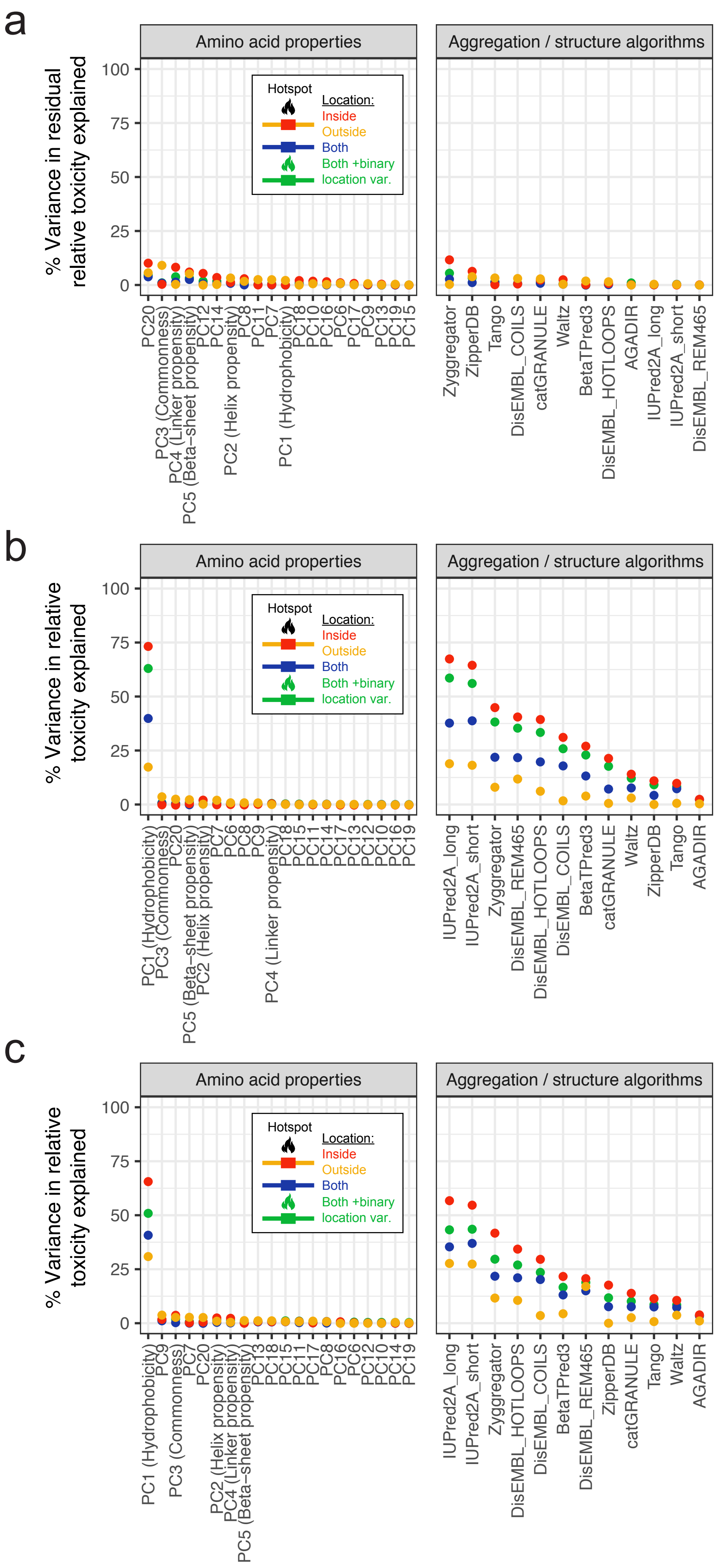


### Extended Data Figure 4

a

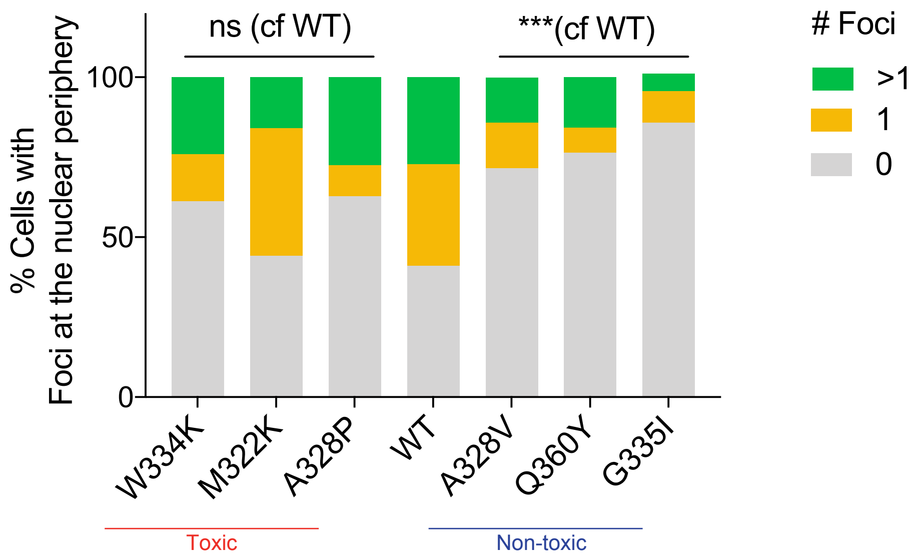


b

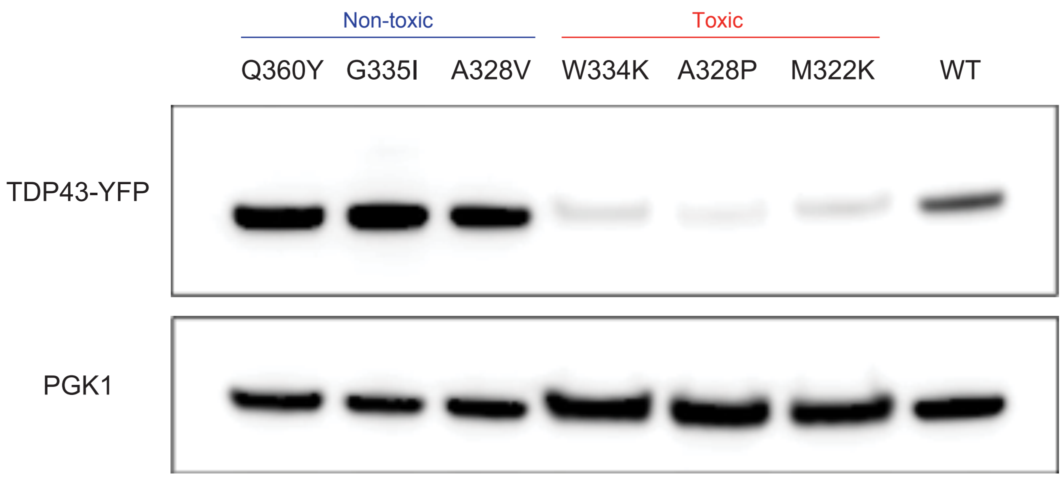


Extended Data Figure 5

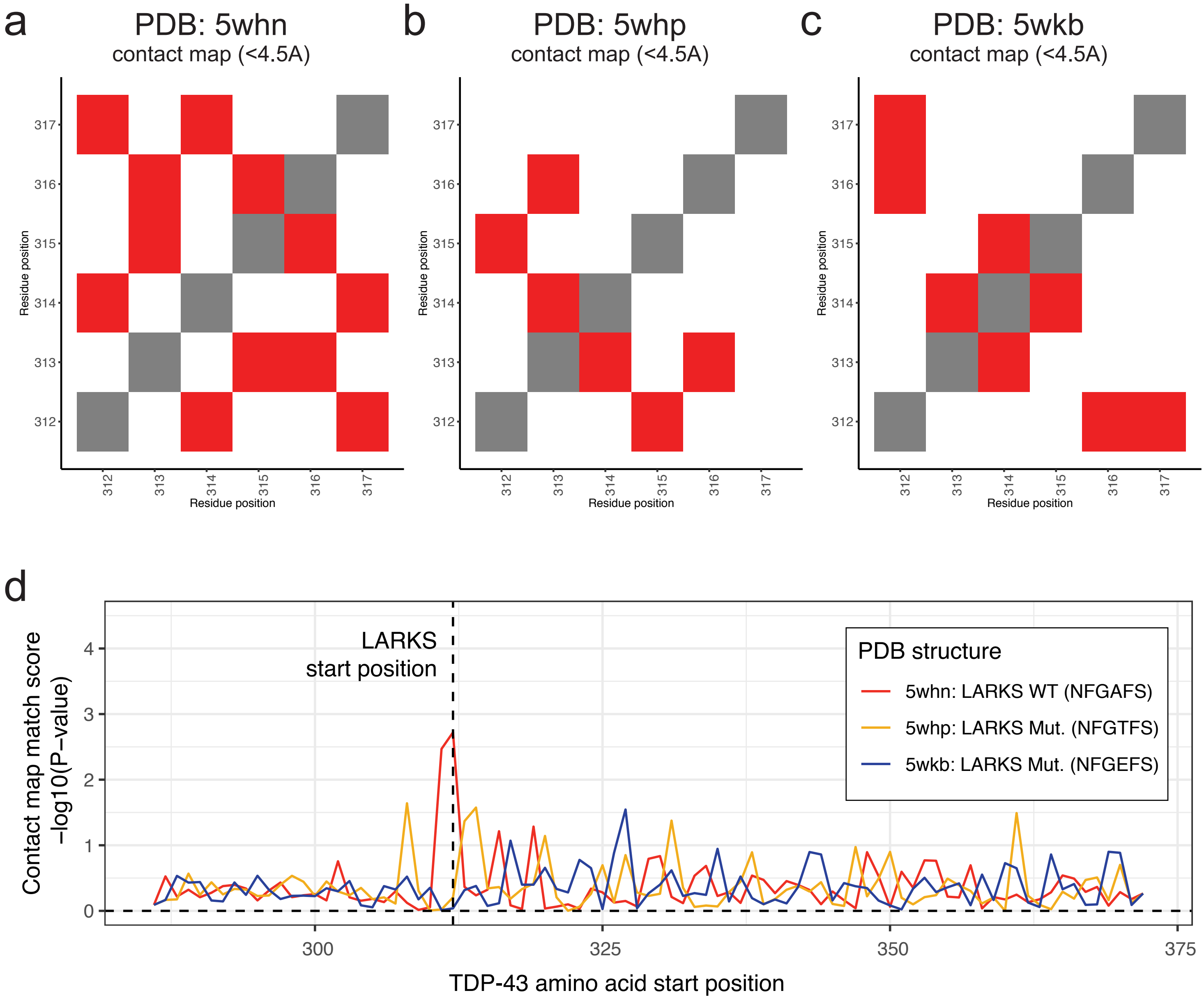


Extended Data Figure 6

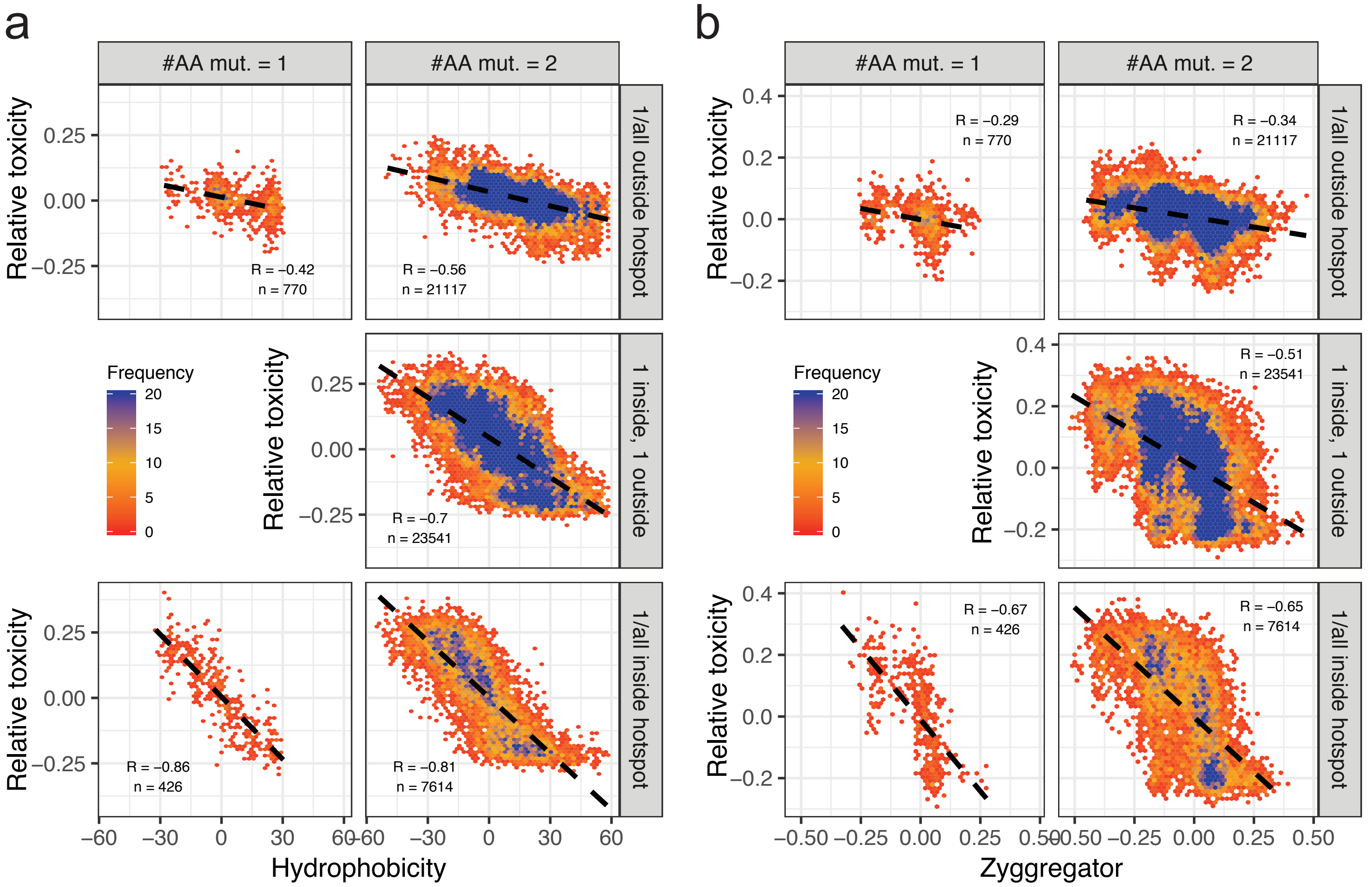


Extended Data Figure 7
